# Epigenetic selection and the DNA methylation signatures of adverse prenatal environments

**DOI:** 10.1101/321711

**Authors:** Elmar W Tobi, Joost van den Heuvel, Bas J. Zwaan, L.H. Lumey, Bastiaan T. Heijmans, Tobias Uller

**Affiliations:** Molecular Epidemiology, Department of Biomedical Data Sciences, Leiden University Medical Center, 2300RC Leiden, The Netherlands; Human Nutrition and Health, Wageningen University & Research, 6708WE Wageningen, The Netherlands; Institute for Cell and Molecular Biosciences, Newcastle University, Campus for Ageing and Vitality, NE4 5PL Newcastle Upon Tyne, United Kingdom; Laboratory of Genetics, Wageningen University & Research, P.O. Box 16, 6700 AA Wageningen, The Netherlands; Department of Epidemiology, Mailman School of Public Health, Columbia University Medical Center, New York, NY 10032, The United States of America; Edward Grey Institute, Department of Zoology, University of Oxford, South Parks Rd, OX1 3PS, Oxford, UK; Department of Biology, University of Lund, Sölvegatan 37, 22362 Lund, Sweden

**Keywords:** DNA methylation, selection, plasticity, developmental origins, DOHAD

## Abstract

Maternal adversity is associated with long-term physiological changes in offspring. These are believed to be mediated through epigenetic mechanisms, including DNA methylation (DNAm). Changes in DNAm are often interpreted as damage or as part of plastic responses of the embryo. We propose that selection on stochastic DNAm differences generated during epigenetic reprogramming after fertilization contributes to the effects of maternal adversity on DNAm. Using a mathematical model of epigenetic reprogramming in the early embryo we predict that this “epigenetic selection” will generate a characteristic reduction in variance of DNAm at selected loci in populations exposed to maternal adversity. We tested this prediction using DNAm data from a human cohort prenatally exposed to the Dutch Famine and confirmed the reduction in DNAm variance, suggesting that epigenetic selection may have occurred. Epigenetic selection should be considered as a possible mechanism linking adversity in pregnancy to offspring health and may have implications for the likely effectiveness of intervention strategies.

## Introduction

Human and animal studies show intriguing associations between adverse events *in utero* and late-life physiology, behavior, and life history (Barker 2007; Brakefield et al. 2005; Lumey et al. 2011). These outcomes may be accompanied by changes in DNA methylation (DNAm) (Heijmans et al. 2009), a key epigenetic mark associated with variation in gene expression that can persist through mitotic divisions (Jaenisch and Bird 2003). Indeed, there is substantial evidence for associations between maternal (mal)nutrition (Joubert et al. 2016; Tobi et al. 2015), glucose metabolism (Quilter et al. 2014) during early gestation and the DNAm profiles of their offspring. However, the mechanism underlying these associations are still poorly understood. Two alternatives are frequently considered (Heijmans et al. 2009). First, adverse conditions in utero may compromise the establishment and maintenance of DNAm (Sharp et al. 2017). Second, DNAm may be actively modified in the embryo to enable a match between the phenotype of the offspring and its current or future environment (e.g. adaptive or predictive developmental plasticity) (Gluckman et al. 2007). If environments do not match expectations, plasticity can cause maladaptation, and this is commonly invoked to explain the increase in non-communicable disease in many parts of the world (Gluckman et al. 2009).

Another possibility that has not yet been fully explored is that embryos with particular epigenetic profiles have different survival probabilities and, as a consequence, there may be selection on epigenetic variation *in utero* (Tobi et al. 2014). Selection on unbiased epigenetic variation (‘epigenetic selection’) is possible if three key conditions hold: (A) there is stochastic variation in DNAm among embryos, (B) specific DNAm patterns confer differential prospects of embryo survival under adverse conditions, and (C) DNAm patterns arising early in life can be transmitted during cell division and persist during development and, in some cases, even into adulthood. There is increasing evidence in the literature that these conditions are met.

Single cell transcriptome and methylome studies have revealed stochastic variation in both gene expression and DNAm between genetically identical cells in common environments (Angermueller et al. 2016). Stochastic genome-wide differences in DNAm may arise during early development as the mammalian genome is stripped of its DNAm after fertilization, followed by a period of global re-methylation in the post-implantation embryo (Messerschmidt et al. 2014). Transient gene expression changes (Greenberg et al. 2016) and regulatory circuits controlled by transcription factors play a crucial role in establishing DNAm patterns (Eckersley-Maslin et al. 2016; Guo et al. 2015), for example, because the binding of transcription factors to promoter and enhancer regions can locally decrease the probability that cytosines become methylated (Bonder et al. 2017; Gebhard et al. 2010; Maurano et al. 2015).

Successful embryonic implantation and further development requires the expression of specific genes to match the blastocyst physiology with the conditions in the uterus and the endometrium (Altmäe et al. 2012). Such genes are associated with transcription regulation, cell adhesion, and signal transduction in the pre-implantation stage, and with cell growth and signaling pathways. The extent to which these processes are aligned is expected to vary between embryos. This will create variation in the likelihood of successful implantation and development. For instance, the success rate of assisted reproduction is linked to the degree of global re-methylation of the early blastocyst placed *in utero* (Li et al. 2017a). Indeed, in humans, more than half of all embryos fail to implant or abort soon after implantation (Wilcox et al. 1999). This number can be substantially higher under stressful conditions (Stein and Susser 1975), as maternal adversity reduces implantation success (Bellver et al. 2013; Dechanet et al. 2011; Li et al. 2017b). We therefore expect that maternal adversity could generate intense selection on gene expression and the associated epigenetic profiles, in particular during implantation.

The DNA methylation maintenance machinery enables transmission of established DNA methylation profiles during cell division (Jaenisch and Bird 2003) despite that DNA methylation is continuously remodeled during fetal development and cell differentiation (Slieker et al. 2015). DNAm patterns established early in development may be maintained (Dominguez-Salas et al. 2014) if these regions do not require extensive remodeling later on. Moreover, the consistent DNAm patterns found in populations that developed under adverse conditions indicate that some of these patterns can persist through adulthood (Heijmans et al. 2008).

Because epigenetic selection removes particular DNAm variants from the population, it may change the observed distribution of DNAm in survivors. Thus, it should be possible to detect signatures of epigenetic selection *in utero* from DNAm data in human populations (similarly to the rationale for detecting signatures of selection in DNA sequence data (Masel and Promislow 2016)). With the aim to derive testable predictions we implemented a mechanistic model that captures the essentials of the dynamics between transcription factor binding and re-methylation (Chen et al. 2013) and extended this model to encompass multiple cells in a whole embryo. We simulated both the plasticity and epigenetic selection scenarios to determine whether epigenetic selection would lead to distinct DNAm signatures at the population level. In particular, we predicted that epigenetic selection, but not plasticity, should result in a reduction in DNAm variance at CpG sites that confer a selective advantage. Finally, to evaluate whether selection on stochastic epigenetic variation has predictive power, and thus warrants further exploration, we compared the predictions from these simulations to empirical DNAm data from individuals exposed to the Dutch Famine during gestation and prenatally unexposed (sibling) controls (Tobi et al. 2015). The Dutch Famine was a severe war-time famine at the end of World War II during which the number of births dropped to almost 50% of normal levels (Susser and Stein 1994), due to a combination of reduced conception, lower implantation success, and an increased rate of fetal deaths in the famine exposed population (Stein and Susser 1975)

## Results

### Modeling dynamic TFBS methylation during development

To generate testable predictions we modified the mechanistic model of Chen *et al*. (Chen et al. 2013) to model the relationship between transcription factor binding and re-methylation. Although this model excludes significant biological details, these details matter primarily insofar as they contribute to consistent effects of maternal adversity on transcriptional regulation (i.e., plasticity) or embryo survival (i.e., selection), as these are the processes that generate differences in DNAm among exposed and unexposed groups. We therefore omit several molecular intermediates in the DNAm machinery and other epigenetic factors. Instead, we focus on the variation in DNAm caused by the inherent stochasticity in DNAm and transcription factor binding, transcription factor concentration, and the effects of gene expression on implantation success. In the Supplementary Information we explore several versions of the basic model we present below, which show the robustness of the model predictions across mechanistic details and parameter space. The full details of the model can be found in the methods and here we only provide a brief and non-technical summary of the model and robustness analyses.

Each of our simulations started with 100 embryos, and every embryo consisted of 50 cells containing 75 independent genes with one transcription binding site (TFBS) each. Thus, we reduced the complexity of gene regulation (i.e., multiple different TFBS and the correlation in methylation patterns across loci (Talens et al. 2010; Teschendorff et al. 2014)), to a representation by one TFBS per gene. We assumed ten CpG dinucleotides (CpGs) per TFBS, but the actual number does not impact on the qualitative behavior of the model (*data not shown*). Since we were interested in the consequences of purely stochastic variation in DNAm, we incorporated no genetic variation between different embryos and no sequence variation between different TFBS.

Our model starts at pre-implantation, when the genome is demethylated and none of the CpGs in the TFBS are methylated. We modeled the dynamics of re-methylation of CpGs within the TFBS as a stochastic process that depends on transcription factor (TF) concentration (Fig. 1A). We assumed equal concentrations of a particular TF in all 50 cells constituting an embryo *in silico*. When a TFBS is occupied by a TF the likelihood that a CpG in a cell will become methylated is decreased (Domcke et al. 2015), as TF binding blocks access for the methylation machinery (Bonder et al. 2017). This process is modelled by simply setting a probability for re-methylation when a TFBS is temporarily vacant. In addition, methylation of a CpG inhibits TF binding (Yin et al. 2017) and hence the likelihood that a neighboring CpG becomes methylated increases. The duration a TFBS is bound by a TF corresponds with the level of gene expression. Since these processes follow simple Michaelis-Menten kinetics the binding of TF and the methylation machinery to a TFBS are inherently stochastic. As a result of these dynamic kinetics both methylation frequencies and gene expression levels are variable between individual cells and embryos (Fig. 1A, Supplemental Fig. S1). We chose to only incorporate this as an inverse relationship between transcription and DNAm as more complex relationships (Bonder et al. 2017), such as including the positive correlation between gene expression and gene body DNA methylation, are unlikely to alter the qualitative behavior of the model (Supplemental information, Supplemental Fig. S2). Finally, we assume that DNAm is heritable during cell division once re-methylation is completed, and, hence, any differences in DNAm established early in life will be detectable in post-natal life.

**Figure 1.**
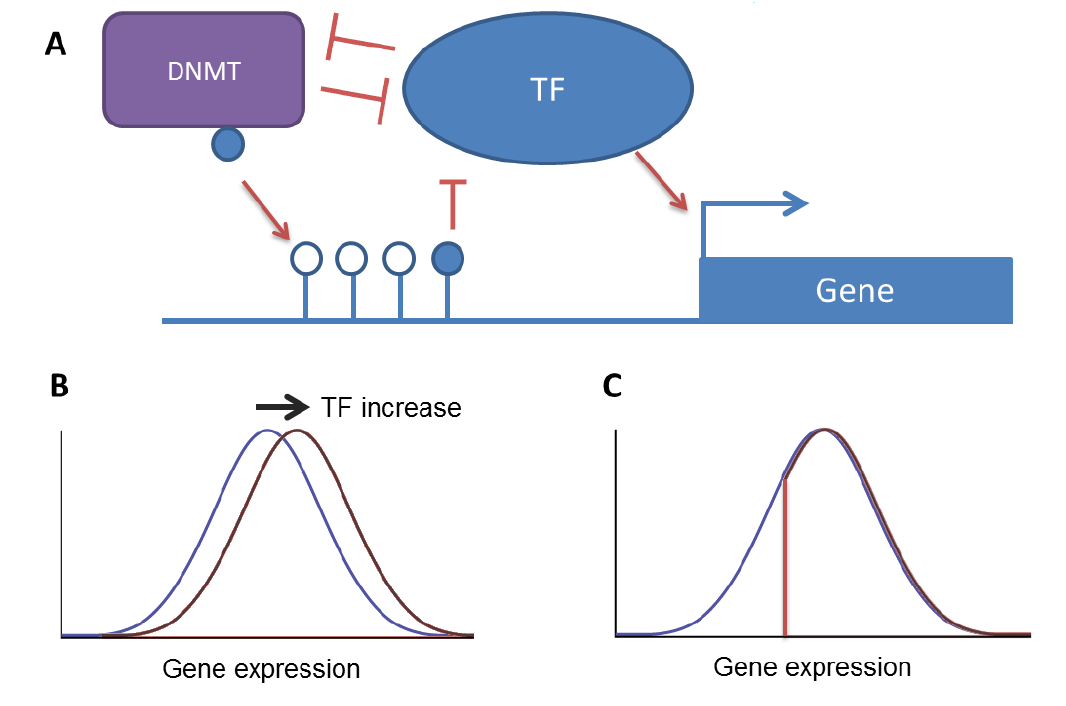
Model of re-methylation and simulations of plasticity and selection. **A)** DNA methylation dynamics during re-methylation in early embryos. Methylation agents such as DNMTs compete with a transcription factor (TF) for the binding at the transcription factor binding site (TFBS). The probability that a TF binds depends on its concentration and the methylation status of the TFBS. DNAm reduces the probability that TF binds and therefore increases the probability that DNMTs methylate a previously unmethylated site in the TFBS. Gene expression is a function of the binding of TF, and is therefore inhibited by DNAm. Once a site is methylated, it has a high probability of remaining methylated during the re-methylation period because of DNAm maintenance. Because the binding of DNMT and TF are all partly stochastic processes, both methylation frequencies and gene expression levels are variable between individuals. **B)** Under the plasticity scenario, we simulate an increase (or decrease, depending on the gene) of [TF] in cases which results in an increase in gene expression (and decreased probability that the linked TFBS becomes methylated). However, variation in gene expression (and thus DNAm) still exists between individuals due to the stochastic nature of the re-methylation dynamics. **C)** Under the scenario of epigenetic selection, we assume that only embryos with a gene expression level higher than some cut-off value (i.e., right of the vertical line) survive selection. As in the plasticity scenario, this produces a shift in mean gene expression. The mean and variation in gene expression and methylation frequencies acan be quantified and compared between the two scenarios.

To generate distributions of DNAm in samples of individuals, we first calculated the mean DNAm for each individual CpG within a TFBS over the 50 cells (each of which are either 0%, 50% or 100% methylated) of an individual embryo at the end of each simulation. We then quantified the mean and variance in methylation frequency for each single CpG between embryos. TF binding is positively related to gene expression of the target gene, which is also quantified as the average gene expression of an individual embryo by taking the mean over all cells. The default state is a simulation without maternal adversity (Figure 2A). Under these assumptions, the modelled distributions of DNAm and the relationship between mean and variance of methylation frequency reproduce empirical patterns of DNAm as measured by genome-wide DNAm arrays in whole blood from the Dutch Hunger Winter Families Study (Lumey et al. 2007) (Figure 2B). Genome-wide DNA patterns are characterized by a majority of CpGs with methylation levels close to 0% and 100% with little to no variation in the population and a minority of intermediately methylated CpGs which show a peak in variation around 50% methylated, resulting in a mean-variance structure following a negative parabola (the higher levels of hypo- and hyper-methylated CpGs in the empirical data is because of the inclusion of CpG islands, which show no to little variation in the population and were therefore not modelled).

**Figure 2.**
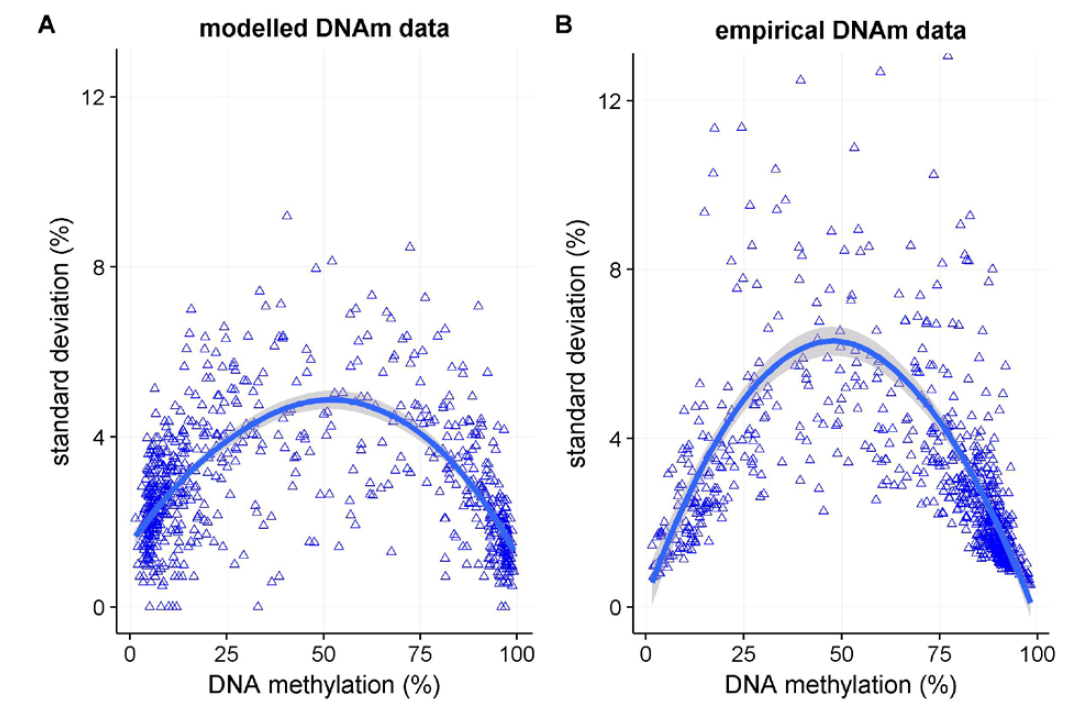
Comparison of modelled and empirical DNAm data. **A)** DNAm for 750 simulated CpGs within 75 TFBS are shown, **B)** DNAm for 750 random CpGs from 463 controls from the Dutch Hunger Winter Families Study are shown. Both panels show the trend (LOESS with span=1.0 in blue and the 95% C.I. in gray, as calculated within the R *ggplots2* package) between mean methylation (x-axis) and standard deviation (y-axis).

### Simulating plasticity and epigenetic selection

We used the above model of re-methylation during development to generate predictions for DNAm patterns under plasticity and epigenetic selection in a series of new simulations with the aim to find a DNAm signature of epigenetic selection in cohorts exposed to maternal adversity. Under plasticity, transcription factor concentration [TF] responds in a coordinated adaptive response to the nutritional conditions *in utero* (Fig. 1B). We therefore modeled plasticity by increasing or decreasing [TF] upon maternal adversity and studied its effects on the DNAm of the corresponding TFBS. In contrast, under epigenetic selection there are no systematic differences in [TF] *in utero*. Instead, under epigenetic selection maternal adversity reduces overall embryo survival by favoring embryos with particular gene expression levels (i.e., those favorable for successful implantation and survival). That is, under this scenario there is selection of a subset of the stochastically arising DNAm profiles (Fig. 1C).

For each simulated scenario, we modelled 150 genes, 75 ‘target genes’ that either respond plastically (plasticity model) or influence the probability of survival (epigenetic selection model), and 75 ‘control genes’, in a population of embryos with or without maternal adversity. At the end of each simulation, when the *in silico* ‘genome’ has completed its re-methylation (i.e., the DNAm levels remain stable), we characterized the resulting DNAm patterns in each group of embryos. Because the number of embryos is reduced by selection, and this may influence the comparison to the plasticity scenario, we started these simulations with a larger number of embryos and used random subsampling to maintain the same population size across all comparisons.

### Model predictions for epigenetic selection

Both plasticity and epigenetic selection caused a shift in mean methylation at the individual CpGs linked to the 75 target genes with a role in plasticity or survival (both increases and decreases in mean DNAm were observed, *data not shown*). The model simulations showed that, under plasticity, the relationship between the mean and variance in methylation was the same in groups of exposed and unexposed individuals (Figure 3A). In contrast, epigenetic selection reduced the variance in DNAm at TFBS that contribute to survival. This reduction is modest at single CpGs, but evident across the entire range of mean DNAm levels (Figure 3B, Supplemental Figure S3). The relationship between the mean and variance for control genes (i.e., genes whose expression does not contribute to embryo survival) was unaffected by selection (Figure 2C). As expected, the disparity in DNAm profiles between embryos surviving under adverse conditions and controls increased with the intensity of selection, that is, when fewer individual embryos survived (Figure 2D, see also Supplemental Figure S3). Hence, the model singles out a modest, but consistent, reduction in variance in DNAm across its entire range (e.g. from 0-100%) as a key feature of epigenetic selection.

**Figure 3.**
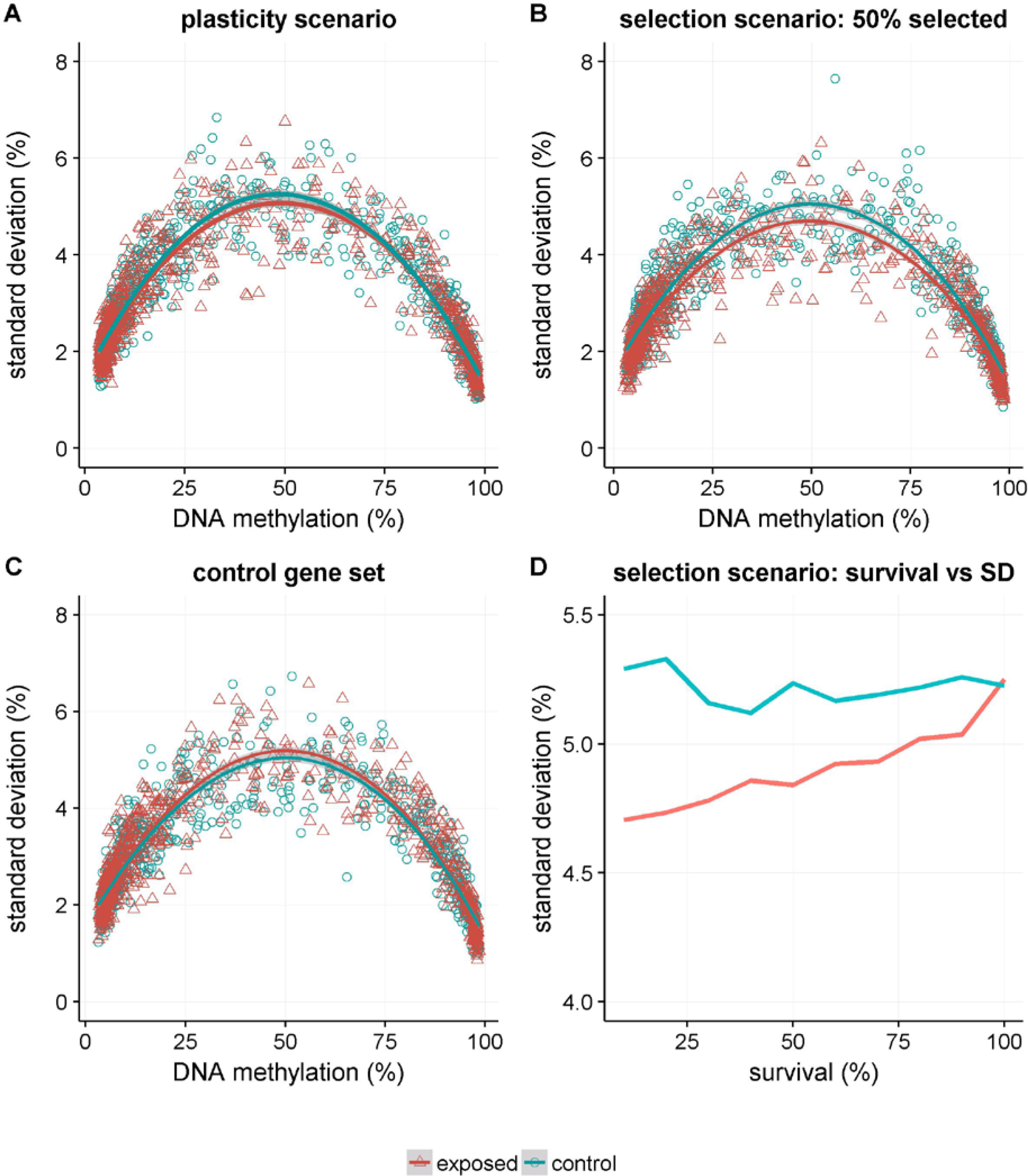
Predictions for the relationship between mean and variance in DNAm. Model predictions for the relationship between mean and standard deviation in DNAm. In each panel the blue circles and/or solid blue trend lines (a LOESS with span=1.0 and 95% C.I. in gray, as calculated by the R *ggplots2* package) denote the observations in the simulated embryos that do no experience prenatal adversity (“controls”). The red pyramids and/or solid red trend lines (identical LOESS settings) are the observations in simulated embryos exposed to maternal adversity (“exposed”). **A)** Relation between mean DNAm and standard deviation (SD) for 750 CpGs in 75 TFBS linked to genes active in the plastic response. **B)** Relation between mean DNAm and standard deviation (SD) for the 750 CpGs in 75 genes linked to survival when 50% of the exposed embryos fail to thrive as a result of the exposure. **C)** The same selection scenario as in panel B, but now for the 750 CpGs in the 75 TFBS linked to genes not vital for survival, i.e., the control gene set. **D)** Standard deviation of methylation under different cut-offs for survival.

### Robustness Analyses

To study the robustness of the variance reduction under epigenetic selection, we performed simulations under different parameter setting, as well as for models with additional molecular detail or alternative assumptions. In a large parameter and scenario space, variance was reduced under selection compared to control conditions (Supplemental Information). Most importantly, the results of the model were robust also under conditions causing lower DNAm maintenance and re-methylation, for instance through a shortage of essential (micro-)nutrients for the methylation machinery. Indeed, only at extreme parameter values for methylation maintenance does the introduced variation in DNAm upon re-methylation becomes sufficiently large to make the effect of epigenetic selection on the mean-variance relationship undetectable (Supplemental Fig. S4). The predicted effect of epigenetic selection on variance reduction was also robust when other relationships between TF binding and the methylation machinery, e.g. when different proportions of TFBSs with TFs recruiting or inhibiting *de novo* methylation, are modelled (Supplemental Fig. S2). These results were obtained from simulations in which all cells had equal concentrations of TF within an embryo. However, in reality a gradient of TF concentration exists for many important developmental TFs. Such a gradient has no considerable effect on variance reduction when epigenetic selection is acting (Supplemental Figure S5). Despite this robustness, it is important to highlight that due the stochastic nature of the modelled re-methylation and selection process itself not all individual simulations show a decrease in mean-variance, nor are they expected to.

### Evaluating model predictions in empirical data

To illustrate how to test for this reduction in variance in empirical DNAm data, and to look for support for epigenetic selection in a human population, we used genome-wide DNAm data in whole blood for 422 individuals prenatally exposed to the Dutch Famine and 463 unexposed (sibling) controls (Supplemental Table 1). The Dutch Famine is considered a quasi-experimental setting and we previously reported on DNAm patterns associated with prenatal famine exposure (Tobi et al. 2014, 2015, 2009), identifying early gestation as a particular sensitive developmental time-frame.

**Table 1.** Difference in variation for differentially methylated CpGs between exposed and controls.

|  | Famine associated CpGs |  |  |  |  | 1K unaffected CpGs |  |
| --- | --- | --- | --- | --- | --- | --- | --- |
| | N probes | ΔSD(%) | Corr<sup>1</sup><br>ΔSD(%) | P 100K permutations<sup>2</sup> | P FDR | Corr<sup>3</sup><br>ΔSD(%) | P 100K permutations<sup>2</sup> |
| Any famine exposure | 543 | -0.1 | 0.0 | 0.32 | 0.32 | 0.01 | 0.74 |
| Exposure period specific |  |  |  |  |  |  |  |
| -10 - 0 | 404 | -0.2 | -0.2 | 0.010 | 0.031 | 0.0 | 0.67 |
| weeks 1-10 | 669 | -0.4 | -0.2 | 0.21 | 0.32 | 0.1 | 0.58 |
| weeks 11-20 | 520 | -0.4 | -0.3 | 0.064 | 0.13 | 0.1 | 0.60 |
| weeks 21-30 | 506 | -0.3 | -0.3 | $7.3 \times 10^{-3}$ | 0.031 | 0.0 | 0.52 |
| weeks 31-delivery | 622 | -0.2 | -0.1 | 0.29 | 0.32 | 0.0 | 0.47 |

1. The mean difference in SD between exposed and controls across CpGs associated with prenatal famine exposure; corrected for age, sex, technical variation and cellular heterogeneity. 2. P value attained by 100K permutations on Beta values corrected for age, sex, technical variation and cellular heterogeneity. 3. The mean difference in SD between exposed and controls across 1K probes not associated with prenatal famine exposure; corrected for age, sex, technical variation and cellular heterogeneity.

We compared the variance in DNAm in prenatally exposed individuals and controls across all individual CpGs putatively associated with famine exposure (at P<0.001) during 10-week time-frames, or during any of these gestational time-frames (“any famine exposure”). These loci are enriched for CpGs with intermediate levels of DNAm at (developmental) enhancers and promoters devoid of CpG islands (Tobi et al. 2014). These famine-associated CpGs were contrasted to 1000 randomly selected control CpGs that had a similar mean and variance as the famine-associated CpGs in the prenatally unexposed, but that were not associated with prenatal famine exposure (P>0.2). Selecting a set of CpGs with the same mean-variance structure ensures that the comparison is not confounded by differences in the mean and variance of the selected subset itself.

The variance in DNA methylation at famine-associated CpGs was significantly lower in individuals whose mothers were exposed to the height of the famine in the 10 weeks before conception (ΔSD=-0.2%, P_FDR_=0.031, Figure 4A). A similar reduction was found in individuals exposed during weeks 21-30 of gestation (ΔSD=-0.3%, P_FDR_=0.031, Figure 4B), but not for the other 10-week time frames, nor when all time frames were combined (Table 1). Both sets of ‘control’ CpGs showed no difference in variance (Table 1). Similar results were obtained when the exposed individuals were compared to their same-sex, unexposed sibling controls (Supplemental Table S2). The lower variance in DNAm in exposed individuals is similar to the model predictions under epigenetic selection.

**Figure 4.**
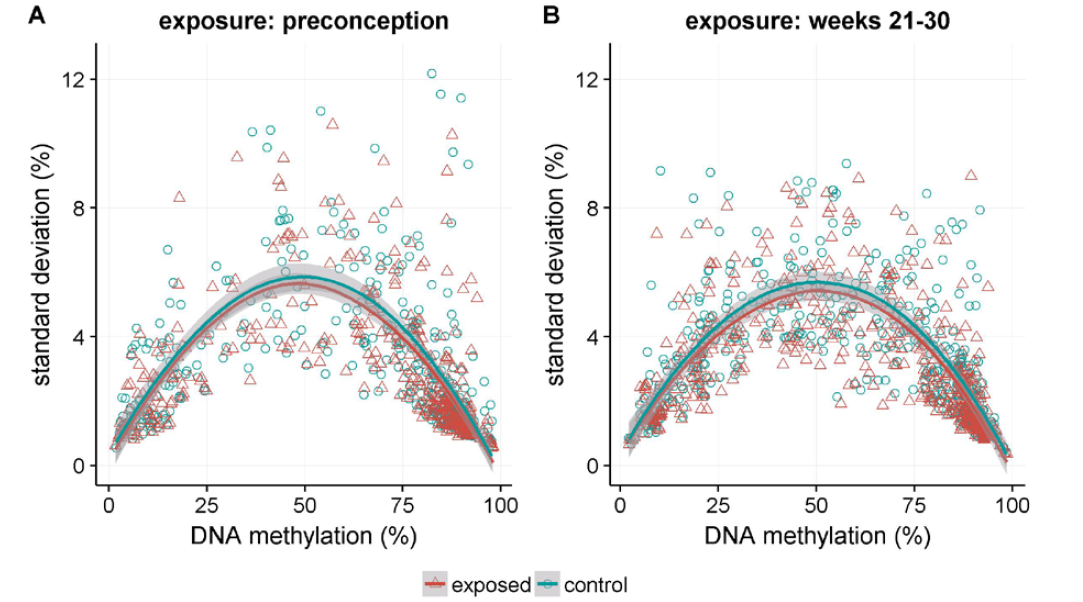
Empirical DNAm data. Empirical data for the relationship between means and standard deviation of DNAm data in the Dutch Hunger Winter Families Study, for **A)** the 404 CpGs associated with preconception famine exposure and **B)**, the 506 CpGs associated with famine exposure during weeks 21-30 of gestation and controls. The mean DNAm (x-axis) and standard deviation (y-axis) and trend between these two variables is given in blue for controls and red for exposed individuals (LOESS with span=1.0 and 95% C.I. in gray, as calculated by the R *ggplots2* package).

## Discussion

Our main aims in this paper were to (i) examine epigenetic selection as a possible explanation for the observed associations between prenatal adversity and DNAm in children and adults, (ii) use a basic mechanistic model in combination with individual-based simulations to identify predictions for the effect of epigenetic selection on DNAm patterns in exposed and control populations, and (iii) demonstrate how to test these predictions on empirical DNAm data and provide a first assessment of the merits of the epigenetic selection hypothesis. The model showed that epigenetic selection reduces the observed DNAm variance of DNAm in exposed compared to unexposed populations. The variance in DNAm may therefore be used as an indicator of epigenetic selection. A first test of this hypothesis suggests that epigenetic selection may indeed be a tenable explanation for the DNAm patterns observed in individuals exposed to maternal adversity during the earliest stages of development *in utero*.

Epigenetic selection is particularly important to consider when prenatal adversity occurs shortly after conception. Epigenetic selection is a plausible process in very early gestation because mortality during human embryonic development is highest directly before and after implantation (Macklon et al. 2002; Wilcox et al. 1999). This period is largely captured in the Dutch Hunger Winter Families study by those pregnancies exposure to famine for at least 10 continues weeks before conception. These individuals were conceived at the height of the famine, when rations fell to 500 kcal/day, and the DNAm variance reduction in this exposed group was indeed observed. There was a similar variance reduction for individuals exposed anytime during the first two trimesters, although this was statistically significant only for gestational weeks 21-30. Overall, these preliminary results indicate that the DNAm pattern associated with early gestational adversity may have been generated by the selective survival of embryos with particular DNAm profiles rather than by embryo plasticity.

However, detecting a signature of epigenetic selection does not rule out that DNAm in embryos could also respond directly to maternal adversity; both processes may occur simultaneously. Plasticity may be a better explanation for mean changes in DNAm that occur late in gestation, when mortality is lower, as it could aid the fetus to better cope with the adverse intrauterine environment. Furthermore, environmental cues become more relevant for predicting postnatal conditions closer to birth (Nettle et al. 2013). An interesting example to explore may be the well-established link between growth in late gestation, reduced nephron number and its associated increased risk for hypertension (He et al. 2010), which may have an epigenetic component (Bogdarina et al. 2010). However, directly testing the adaptive plasticity hypothesis in humans is difficult (Brakefield et al. 2005), since reliable estimates on Darwinian fitness are difficult to obtain in modern societies. Ruling out epigenetic selection may therefore be especially important to studies that invoke adaptive plasticity to explain associations between maternal adversity, DNAm, and phenotypic characters.

Our test for a signature of epigenetic selection in empirical data of the Dutch Famine is only a first step to investigate the scope for the epigenetic selection hypothesis. More detailed studies are needed, in particular since we were only able to measure the DNAm patterns in blood six decades after the exposure. Furthermore, only moderate differences in variance were observed in the quasi-experimental setting of the Dutch Famine, despite the severe nutritional deprivation. Moreover, our simulations suggest that the stochastic nature of re-methylation means that large data sets are required to robustly identify regions with a lower-than-expected DNAm variance. The recent formation of large consortia with detailed information on contemporary prenatal conditions and genome-scale DNAm data holds great potential in this respect (Felix et al. 2017). Experimental studies in animal models is another promising route to provide more direct tests for epigenetic selection.

From a theoretical perspective, it will be important to also study how different factors such as age and tissue heterogeneity, DNA sequence variation and the developmental process itself (e.g., canalization (Pujadas and Feinberg 2012)) can affect the variance of DNAm. In particular, a reduced variance could reflect a more canalized regulation of gene expression. Although the key predictions were unaffected by a number of mechanistic detail regarding the process of re-methylation, including that plasticity is caused by shortage of substrate or co-factors (e.g. a lower rate of re-methylation and maintenance), it is important to note that the magnitude of the reduction in variance can be affected. Indeed, other forms of selection (e.g., increased survival of both extremes of a distribution) may increase the variance within a population. Another possibility is to impose selection within, rather than between, blastocysts. Selection of particular cells within an embryo occurs during normal development (Sancho et al. 2013) and may likewise be hypothesized to play a role in fetal programming. Future modelling efforts should also strife to incorporate new emerging insights on early human embryonic histone dynamics, as histone modifications may offer another molecular signature to test for epigenetic selection. Finally, although we make no assumptions about whether or not epigenetic selection *per se* is adaptive, it may be interesting to explore the conditions under which it evolves under natural selection (Nettle et al. 2013; van den Heuvel et al. 2016).

Understanding the mechanisms of DNAm changes in an adverse prenatal environment has practical and clinical implications. Under epigenetic selection, aberrant DNAm profiles of survivors are not caused by adversity during early life; they arise from the effects of stochastic variation in DNAm on the odds of implantation or post-implantation survival. This hypothesis makes no assumption about specific adaptive or maladaptive consequences of those DNAm profiles that are observed later in life. By contrast, under adaptive plasticity it is posited that an adverse maternal environment induces specific epigenetic profiles and that such adaptations may increase the risk of disease in a mismatched post-natal environment (Gluckman and Hanson 2006). Hence, while prevention or intervention measures in fetal life could mitigate or neutralize adverse plastic responses, under epigenetic selection such interventions would merely reduce the selection pressure and not necessarily prevent adverse epigenetic profiles that arise stochastically. Thus, the distinction between induction of DNAm profiles *versus* selection of pre-existing DNAm profiles may be important for managing the expectations of intervention programs and the ongoing discussion of (epi)genetic determinism (Holbrook 2015; Waggoner and Uller 2015), including concerns that we are placing blame on past generations for the diseases of our own (Richardson et al. 2014).

In summary, we propose that differential survival of embryos with particular epigenetic profiles (‘epigenetic selection’) is a tenable explanation for the observed late-life differences in DNAm after early prenatal adversity in humans. By formulating and modelling this alternative, we show that it generates predictions that can be tested using data that are available in human populations. Predicted DNAm profiles from epigenetic selection models are in close agreement with empirical findings among men and women with prenatal exposure to the Dutch famine of 1944-45. This implies that epigenetic selection needs further consideration as a biological mechanism by which early life adversity in humans could have long lasting impacts.

## Methods

### Mathematical model

We modelled CpGs (a vector, *M*), which can be in a methylated or un-methylated state. Generally, transcription factors (TFs) and proteins that methylate CpGs compete for binding to the DNA (Domcke et al. 2015) (see Supplemental Fig. S6 for a schematic overview of the model characteristics). Demethylation of target sites leads to an increase in TF binding (Maurano et al. 2015), while the binding of TFs results in resistance to *de novo* DNAm by DNMTs (Brandeis et al. 1994; Maurano et al. 2015). To encapsulate these dynamic behaviours we follow the logic of the modelling approach by Chen *et al*. (Chen et al. 2013). The probability of binding of TF and DNAm is negatively related to the presence of the other. A higher activity of TFs will therefore reduce the probability that DNA will become methylated, while more methylation in a transcription factor binding site (TFBS) decreases the probability of TF binding. DNAm therefore causes a lower gene expression for the target gene. For each TFBS we modelled 10 CpGs and each TFBS was replicated in 50 cells (the approximate maximum number of cells in a human pre-implantation blastocyst (Herbert et al. 1995)). Molecular stochasticity gives rise to variation in DNAm between cells and between individuals. We model no differences in DNA sequence between embryos or TFBS within a cell of one embryo (we did therefore not explicitly model CpG islands, which are hypo or hyper-methylated and generally lack DNAm variation (Jaffe et al. 2012)).

The dynamics of gene expression and DNAm depend on the concentration of transcription factors, which reduces the ability of DNAm transferases (DNMT) to methylate CpGs (i.e., increased availability of TFs reduces methylation on average, Figure S1A). We modelled gene expression (*G*) as,

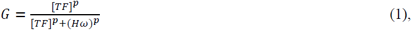

where *[TF]* is the concentration of active transcription factor, the parameter *H* is the half-saturation value, *p* determines the rate at which gene expression increases with *[TF]*, and the parameter *ω* modulates this relationship and is itself a function of methylation state. Following Chen *et al.* (Chen et al. 2013) for a given locus *i* in cell j, *ω*_*ij*_ is

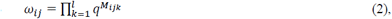

where *l* is the number of modifiable loci (CpGs) in the TFBS, and *q* is the effect of methylation of a single locus on *ω*_*ij*_. This is 1 if methylation has no effect or >1 for a negative effect of methylation on transcription. Parameter *M*_*ijk*_ thus describes the methylation status for each TFBS at the *k*th locus, in the *j*th cell for the *i*th individual. In all simulation that follow we set *q* = 1.2. Figure S2 shows the relationship between transcription factor concentration, *[TF],* and gene expression, G, for different number of CpGs, M.

The dynamics of re-methylation over time is defined by the probability of an un-methylated site becoming methylated and *vice versa*. The probability of methylation of an un-methylated site is inversely related to the binding of a transcription factor, i.e.,

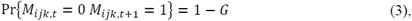

where *M*_*ijk,t*_ and *M*_*ijk,t+1*_ are the states of methylation of the *k*th locus (individual *i* and cell *j*) in a TFBS at times *t* and *t+1* respectively, and *G* is the gene expression (eq. [1], Figure S1B). Therefore, the probability of a locus to remain un-methylated is equal to the concentration of a transcription factor. We assume that the latter is independent of transcription, i.e.,

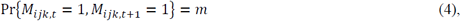

where *m* is a constant (Supplemental Fig. S6C). The probability that a CpG site loses its methylation is therefore (1-*m*).

We initiate every simulation with a transcription factor concentration and all loci begin as un-methylated, i.e. all values of *M* are set to zero. We simulated ‘genes’ with different levels of expression by stepwise increasing *[TF]* (details in Table S3). Gene expression is calculated per cell (Figure S2) and, within an embryo, *[TF]* is equal for all cells. Because methylation status as determined by eq. [2] is zero for all potential methylation sites, the *[TF]* determines the probability of CpG dinucleotides being methylated, following eq. [3]. Then for every given site we compare a randomly drawn number from a uniform distribution with the outcome of eq. [3]. A CpG dinucleotide is methylated when this random number is lower than the probability calculated under eq. [3]. In the next time step, some sites will be methylated, while others remain un-methylated. Again by eq. [2] we determine the methylation status of every site and determine the probability of methylation for sites that are not methylated. However, for sites that are methylated we determine the probability that this methylation is maintained, according to eq. [4]. Therefore, for every time step we iterate over all individuals, cells and TFBS and CpGs within TFBS and determine states and state changes. For every time step we can calculate the gene expression of an individual by taking the mean over all cells (Supplemental Fig. S7). Furthermore, for every CpG site we can do the same (Supplemental Fig. S8). Note that for a specific gene we therefore generate one gene expression level, while multiple mean levels of methylation exist (compare Supplemental Fig. S7 with S8).

### Simulations

For the epigenetic selection hypothesis, the likelihood of implantation of an embryo depends on its gene expression level, which itself is not caused by maternal adversity but varies as a result of the stochastic nature of TF and DNAm binding (as outlined above). The resulting gene expression levels, as well as the values of methylation frequencies (mean methylation over all cells) are saved before (as control) and after selection (as cases). The intensity of selection was varied through the proportion of embryos that would successfully implant. Because the number of individuals after selection is lower, and this could potentially influence the variation, we subsample the controls to the same *N* as the cases. To induce variation in the mean methylation between different genes we altered the *[TF]*. Thus, the effects of selection are compared between controls and selected embryos for a distribution of gene expression varying on a continuous scale between 0 and 1 in mean methylation frequency (see Figure 1).

For the adaptive plasticity hypothesis, cases (those under adversity) had a simulated concerted change in *[TF]* for a gene responding to maternal adversity (as a simulation of adaptive plasticity). On the contrary, to simulate control individuals, *[TF]* did not change (in effect creating a mean difference in gene expression between cases and controls). We assumed similar *[TF]* for all cells within individual embryo’s and furthermore assumed no differences in *[TF]* between individuals at the start of the simulation. As above, the mean and standard deviation in methylation frequencies were computed for individuals CpG dinucleotides and mean and standard deviation were compared between controls and cases (see Figure 2A). All values of constants used can be found in Table S3. Means and standard deviations were calculated in the in the R programming environment (R Development Core Team, 2009), while all simulations were performed in C++.

### Study subjects

The Dutch Hunger Winter Families study is described in detail elsewhere (Lumey et al. 2007). In short, historical birth records were retrieved from three institutions in famine-exposed cities in the Western Netherlands. In total 2,417 singleton births between 1 February 1945 and 31 March 1946 of infants whose mothers were exposed to the famine during or immediately preceding that pregnancy were identified. In the birth records of the same institutions we sampled an additional 890 births from mothers not exposed to famine during this pregnancy from 1943 and 1947 to serve as time-controls. An address was obtained for 95% of the individuals (2,300 individuals). These were invited to participate in a telephone interview and in a clinical examination, together with a same-sex sibling not exposed to the famine to serve as a family-control. We have thus collected both family controls of exposed and unexposed individuals. We conducted 1,075 interviews and 971 clinical examinations (of 345 clinic births without a matched sibling and 313 with a matched sibling) between 2003 and 2005. The study was approved by the Institutional Review Board of Columbia University Medical Center and by the Medical Ethical Committee of Leiden University Medical Center and the participants provided verbal consent at the start of the telephone interview and written informed consent at the start of the clinical examination.

### Famine exposure

Food rations were distributed centrally and during the famine and the percentage of calories from proteins, fat, and carbohydrates was constant during the famine period (Burger et al. 1948). Rations were below 900 Kcal/day between November 26, 1944 and May 15, 1945 and we defined famine exposure by the number of weeks during which the mother was exposed to <900 kcal/day after the last menstrual period (LMP) recorded on the birth record (or when missing or implausible [12%], as estimated from the LMP calculated from the birth weights and the date of birth (Lumey et al. 2007). We considered the mother exposed in gestational weeks 1-10, 11-20, 21-30, or 31 to delivery if these gestational time windows were entirely contained within this period and had an average exposure of <900kcal/day during an entire gestation period of 10 weeks. As the famine lasted 6-months some participants were exposed to famine during two adjacent 10-week periods. Pregnancies with LMP between 26 November 1944 and 4 March 1945 were thus considered exposed in weeks 1-10; between 18 September 1944 and 24 December 1944 in weeks 11-20; between 10 July 1944 and 15 October 1944 in weeks 21-30; and between 2 May 1944 and 24 August 1944 in weeks 31 to delivery. We define individuals exposed to one or at most two of these definitions exposed to ‘any’ gestational exposure. This definition does not cover individuals with LMPs between 5 March 1945 and 15 May 1945 who were exposed to extreme famine but for shorter periods. Individuals with a LMP between 1 February and 12 May 1945 were exposed to an average of <900kcal/day for an entire 10 weeks before conception (and up to 8 weeks post-conception) and are denoted as the “preconception” group.

### DNA methylation data

DNAm was measured using the Illumina Infinium Human Methylation 450K BeadChip *(450k array)* and described in detail elsewhere (Tobi et al. 2015). Briefly, samples were randomly distributed ensuring similar distributions of exposure periods, sex ratios and mean ages per 96-well plate and 450K array, keeping sibling pairs together, but randomly assigned to either the left or right column of the 450K array. We assessed data quality using both sample dependent and sample independent quality metrics using the *Bioconductor* package *MethylAid* (van Iterson et al. 2014). Bisulfite conversion efficiency was assessed using dedicated 450K probes and sequencing the *IGF2* DMR0 (Tobi et al. 2012) of a random set of samples. Sample swaps were excluded by re-measuring a subset of the genotypes measured on the 450K array with MASSARRAY and checking the gender of samples using all X-chromosomal CpG dinucleotides. We used *noob* and *Functional Normalization* as implemented in the *minfi* package (Aryee et al. 2014) using six principal components to normalize data. Individual measurements with a detection p-value > 0.01 or zero intensity value in the used color-channel were set as missing. The measurement success rate per sample was >99%. Next, we removed a-specific/polymorphic, non-autosomal, <95% success rate, and completely methylated or unmethylated (in bisulfite sequencing datasets) probes.

### Statistics

We previously used generalized estimation equations (GEE) with a Gaussian link function to evaluate the association between DNAm percentage (the 450k array β-value x 100) and famine exposure, and the analyses here follow this work (Tobi et al. 2015). In short, to identify CpGs associated with famine we ran a separate analysis per 10-week exposure period and for ‘any’ gestational exposure, comparing individuals that meet an exposure definition to all 463 prenatally unexposed time- and family-controls. Using GEE, we controlled for correlation within sibships and adjusted for age, gender, row on the 450k array and technical batch (bisulfite conversion plate and scan batch). We adjusted for cellular heterogeneity of whole blood, the tissue studied here, by incorporating the first three principal variance components as their proxy measure. These PCAs were highly correlated (rho>0.9) with one or multiple cell fractions imputed with the Houseman method and have the benefit that they are not correlated among themselves, which allows the assessment of the level of influence of cell heterogeneity on the outcomes as it eliminates co-linearity in the model. In addition, given that the PCAs are an abstraction (e.g. reflect patterns in the data) there is also the potential that these PCAs will capture variation linked to other unmeasured cell fractions. We took CpG dinucleotides further for an assessment of the variance in DNAm when the analysis for a difference in DNAm between famine exposed and controls showed a p-value of <0.001. As control set we took 1000 CpG dinucleotides not associated with famine exposure (P>0.2) drawn by using de D-optimum criterion, which ensures that the 1000 reflect the distribution of both the mean and variance of the methylation of the famine associated CpG dinucleotides.

For each single exposure period we calculated the difference in variance as follows for both sets of CpGs; we first corrected the Beta values for age, gender, row on the 450k array and technical batch (bisulfite conversion plate and scan batch) and cellular heterogeneity using a simple linear regression. Next, we calculated that standard deviation (SD) (as measure of the variation) of DNAm (%) per CpG for the exposed and controls separately and finally calculated the mean difference between the famine exposed and (sibling) controls over all selected CpGs. The significance of this difference was established by 100,000 permutations as the correlation between CpGs and within sibling pairs and the parabolic relation between SD and mean methylation violates assumptions required for (non)-parametric tests. We performed restricted permutations using the R package *permute* (version 0.9-0) as to assess how remarkable our observation is given our largely paired study design. This entails randomly assigning the 0-1 or 0-0 status for famine exposure status between sibling pairs (we do not have 1-1 pairs) and then randomly flipping exposure status within a pair. In conjuncture, we randomly assigned famine status in the individuals without a sibling in the study. Each permutation had exactly the same number of exposed individuals in the sibling pairs and unrelated individuals as in the actual cohort.

## List of Abbreviations

DNAm: (DNA methylation)
CpG: (cytosine-guanine dincleotide)
SD: (standard deviation)
GEE: (Generalized estimating equation)
PCA: (principal component analysis)
450k array: (the Illumina Infinium Human Methylation 450K BeadChip)
LMP: (last menstrual period)
TF: (transcription factor)
[TF]: (transcription factor concentration)
TFBS: (transcription factor binding site)
DOHAD: (developmental origins of human adult disease).

## Graphical Abstract

Graphical presentation of the plasticity (A1) and epigenetic selection (A2) scenarios according to the model. Under plasticity, gene expression increases as a response to the adverse environment, while under epigenetic selection the gene expression of embryos that are allowed to implant is shifted (indicated by the vertical lines). Both scenarios lead to a shift in mean DNAm (shown as a decrease in DNAm, but for individual CpGs this may be an increase or decrease) in offspring exposed to maternal adversity (B1 and B2). Under the plasticity scenario this leads to an overlapping standard deviation distribution across the natural range in DNAm (C1), while under the epigenetic selection scenario standard deviation is decreased (C2). The middle column represents the Dutch Famine data where exposed show a decrease in late-life health compared to controls. Under adversity, the standard deviation of methylation shows a relative decrease in variance of methylation frequency, consistent with the prediction of the selection scenario.

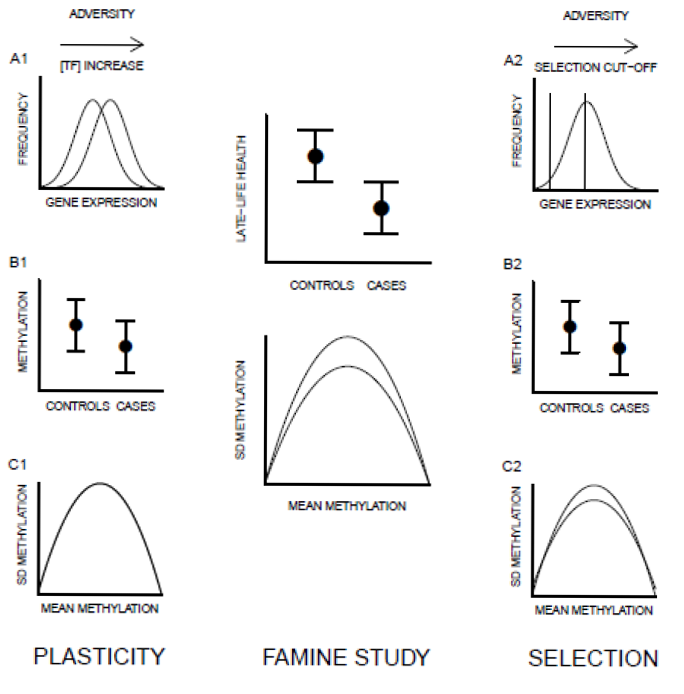

## Ethics approval and consent to participate

The Dutch Hunger Winter Families study was approved by the Institutional Review Board of Columbia University Medical Center and by the Medical Ethical Committee of Leiden University Medical Center and the participants provided verbal consent at the start of the telephone interview and written informed consent at the start of the clinical examination.

## Consent for publication

Not applicable

## Availability of data and materials

All DNAm data generated or analyzed during this study have been published (Tobi et al. 2015), these data are available from the corresponding author (BTH) on reasonable request

## Competing Interests

The authors declare that they have no competing interests.

## Funding

This study was supported by The European Union’s Seventh Framework Program IDEAL [259679], Per-Eric and Ulla Schyberg’s Foundation [140423], and the U.S. National Institutes of Health [R01-HL067914 and R01AG042190]. TU was supported by the Royal Society of London, the John Templeton Foundation [60501], and the Knut and Alice Wallenberg Foundations. EWT was supported by a VENI grant from the Netherlands Organization for Scientific Research [91617128]. The funders had no role in study design, data collection, analysis, decision to publish or preparation of the manuscript.

## Author contributions

Conceptualization: EWT, JvdH, BJZ, BTH, TU. Methodology: EWT, BJZ and JvdH. Investigation: EWT, JvdH and LHL. Formal Analysis: EWT and JvdH. Validation: EWT and JvdH. Software: JvdH. Resources: LHL. Data curation: EWT, LHL. Writing – original draft: EWT, JvdH, BTH, TU. Writing – Review & Editing: LHL, BJZ. Visualization: EWT and JvdH. Supervision: BJZ, BTH and TU. Project administration: LHL, BTH and TU. Funding Acquisition: LHL, BJZ, BTH and TU.

## Acknowledgements

We thank the participants of the EU 7^th^ framework project IDEAL for discussion. We express our gratitude to the participants of the Dutch Famine Families study and the staff of TNO Quality of Life for contact tracing. At the Leiden University Medical Center, we wish to acknowledge the staff of the department of Gerontology and Geriatrics Study Center for the physical examinations and the Central Clinical Chemical Laboratory for extracting DNA.

## References

Altmäe S, Reimand J, Hovatta O, Zhang P, Kere J, Laisk T, et al. 2012. Research resource: interactome of human embryo implantation: identification of gene expression pathways, regulation, and integrated regulatory networks. Mol Endocrinol 26:203–17; doi:10.1210/me.2011-1196.

Angermueller C, Clark SJ, Lee HJ, Macaulay IC, Teng MJ, Hu TX, et al. 2016. Parallel single-cell sequencing links transcriptional and epigenetic heterogeneity. Nat Methods 13:229–232; doi:10.1038/nmeth.3728.

Aryee MJ, Jaffe AE, Corrada-Bravo H, Ladd-Acosta C, Feinberg AP, Hansen KD, et al. 2014. Minfi: a flexible and comprehensive Bioconductor package for the analysis of Infinium DNA methylation microarrays. Bioinformatics 30:1363–1369; doi:10.1093/bioinformatics/btu049.

Barker DJ. 2007. The origins of the developmental origins theory. J Intern Med 261: 412–417.

Bellver J, Pellicer A, Garcia-Velasco JA, Ballesteros A, Remohi J, Meseguer M. 2013. Obesity reduces uterine receptivity: clinical experience from 9,587 first cycles of ovum donation with normal weight donors. Fertil Steril 100:1050–1058; doi:10.1016/j.fertnstert.2013.06.001.

Bogdarina I, Haase A, Langley-Evans S, Clark AJL. 2010. Glucocorticoid effects on the programming of AT1b angiotensin receptor gene methylation and expression in the rat. PLoS One 5:e9237; doi:10.1371/journal.pone.0009237.

Bonder MJ, Luijk R, Zhernakova D V, Moed M, Deelen P, Vermaat M, et al. 2017. Disease variants alter transcription factor levels and methylation of their binding sites. Nat Genet 49:131–138; doi:10.1038/ng.3721.

Brakefield PM, Gems D, Cowen T, Christensen K, Grubeck-Loebenstein B, Keller L, et al. 2005. What are the effects of maternal and pre-adult environments on ageing in humans, and are there lessons from animal models? Mech Ageing Dev 126:431–438; doi:10.1016/j.mad.2004.07.013.

Brandeis M, Frank D, Keshet I, Siegfried Z, Mendelsohn M, Nemes A, et al. 1994. Sp1 elements protect a CpG island from de novo methylation. Nature 371:435–438; doi:10.1038/371435a0.

Burger BCE, Drummond HR, Sandstead JC. 1948. Malnutrition and Starvation in Western Netherlands, September 1944-July 1945. 1st ed. General State Printing Office:The Hague.

Chen C-C, Xiao S, Xie D, Cao X, Song C-X, Wang T, et al. 2013. Understanding variation in transcription factor binding by modeling transcription factor genome-epigenome interactions. PLoS Comput Biol 9:e1003367; doi:10.1371/journal.pcbi.1003367.

Dechanet C, Anahory T, Mathieu Daude JC, Quantin X, Reyftmann L, Hamamah S, et al. 2011. Effects of cigarette smoking on reproduction. Hum Reprod Update 17:76–95; doi:10.1093/humupd/dmq033.

Domcke S, Bardet AF, Adrian Ginno P, Hartl D, Burger L, Schubeler D. 2015. Competition between DNA methylation and transcription factors determines binding of NRF1. Nature 528:575–579; doi:10.1038/nature16462.

Dominguez-Salas P, Moore SE, Baker MS, Bergen AW, Cox SE, Dyer RA, et al. 2014. Maternal nutrition at conception modulates DNA methylation of human metastable epialleles. Nat Commun 5:3746; doi:10.1038/ncomms4746.

Eckersley-Maslin MA, Svensson V, Krueger C, Stubbs TM, Giehr P, Krueger F, et al. 2016. MERVL/Zscan4 Network Activation Results in Transient Genome-wide DNA Demethylation of mESCs. Cell Rep 17:179–192; doi:10.1016/j.celrep.2016.08.087.

Felix JF, Joubert BR, Baccarelli AA, Sharp GC, Almqvist C, Annesi-Maesano I, et al. 2017. Cohort Profile: Pregnancy And Childhood Epigenetics (PACE) Consortium. Int J Epidemiol 16:10–14; doi:10.1093/ije/dyx190.

Gebhard C, Benner C, Ehrich M, Schwarzfischer L, Schilling E, Klug M, et al. 2010. General transcription factor binding at CpG islands in normal cells correlates with resistance to de novo DNA methylation in cancer cells. Cancer Res 70:1398–1407; doi:10.1158/0008-5472.CAN-09-3406.

Gluckman PD, Hanson MA. 2006. The conceptual basis for the developmental origins of health and disease. In: Developmental origins of health and disease. Cambridge University Press:Cambridge. 33–50.

Gluckman PD, Hanson MA, Bateson P, Beedle AS, Law CM, Bhutta ZA, et al. 2009. Towards a new developmental synthesis: adaptive developmental plasticity and human disease. Lancet (London, England) 373:1654–1657; doi:10.1016/S0140-6736(09)60234-8.

Gluckman PD, Hanson MA, Beedle AS. 2007. Early life events and their consequences for later disease: a life history and evolutionary perspective. Am J Hum Biol 19:1–19; doi:10.1002/ajhb.20590.

Greenberg MVC, Glaser J, Borsos MMM, Marjou F El, Walter M, Teissandier AAA, et al. 2016. Transient transcription in the early embryo sets an epigenetic state that programs postnatal growth. Nat Genet 49; doi:10.1038/ng.3718.

Guo F, Yan L, Guo H, Li L, Hu B, Zhao Y, et al. 2015. The transcriptome and DNA methylome landscapes of human primordial germ cells. Cell 161:1437–1452; doi:10.1016/j.cell.2015.05.015.

He K, Zhao H, Wang Q, Pan Y. 2010. A comparative genome analysis of gene expression reveals different regulatory mechanisms between mouse and human embryo pre-implantation development. Reprod Biol Endocrinol 8:41; doi:10.1186/1477-7827-8-41.

Heijmans BT, Tobi EW, Lumey LH, Slagboom PE. 2009. The epigenome: Archive of the prenatal environment. Epigenetics 4:526–531; doi:10.4161/epi.4.8.10265.

Heijmans BT, Tobi EW, Stein AD, Putter H, Blauw GJ, Susser ES, et al. 2008. Persistent epigenetic differences associated with prenatal exposure to famine in humans. Proc Natl Acad Sci U S A 105; doi:10.1073/pnas.0806560105.

Herbert M, Wolstenholme J, Murdoch AP, Butler TJ. 1995. Mitotic activity during preimplantation development of human embryos. J Reprod Fertil 103: 209–214.

Holbrook JD. 2015. An epigenetic escape route. Trends Genet 31:2–4; doi:10.1016/j.tig.2014.09.007.

Jaenisch R, Bird A. 2003. Epigenetic regulation of gene expression: how the genome integrates intrinsic and environmental signals. Nat Genet 33 Suppl:245–254; doi:10.1038/ng1089.

Jaffe AE, Feinberg AP, Irizarry RA, Leek JT. 2012. Significance analysis and statistical dissection of variably methylated regions. Biostatistics 13:166–178; doi:10.1093/biostatistics/kxr013.

Joubert BR, den Dekker HT, Felix JF, Bohlin J, Ligthart S, Beckett E, et al. 2016. Maternal plasma folate impacts differential DNA methylation in an epigenome-wide meta-analysis of newborns. Nat Commun 7:10577; doi:10.1038/ncomms10577.

Li G, Yu Y, Fan Y, Li C, Xu X, Duan J, et al. 2017a. Genome wide abnormal DNA methylome of human blastocyst in assisted reproductive technology. J Genet Genomics 44:475–481; doi:10.1016/j.jgg.2017.09.001.

Li R, Wu J, He J, Wang Y, Liu X, Chen X, et al. 2017b. Mice endometrium receptivity in early pregnancy is impaired by maternal hyperinsulinemia. Mol Med Rep 15:2503–2510; doi:10.3892/mmr.2017.6322.

Lumey LH, Stein AD, Kahn HS, Van der Pal-de Bruin KM, Blauw GJ, Zybert PA, et al. 2007. Cohort profile: the Dutch Hunger Winter families study. Int J Epidemiol 36:1196–1204; doi:10.1093/ije/dym126.

Lumey LH, Stein AD, Susser E. 2011. Prenatal famine and adult health. Annu Rev Public Health 32:237–62; doi:10.1146/annurev-publhealth-031210-101230.

Macklon NS, Geraedts JPM, Fauser BCJM. 2002. Conception to ongoing pregnancy: the “black box” of early pregnancy loss. Hum Reprod Update 8: 333–343.

Masel J, Promislow DEL. 2016. Answering evolutionary questions: A guide for mechanistic biologists. BioEssays 38:704–711; doi:10.1002/bies.201600029.

Maurano MT, Wang H, John S, Shafer A, Canfield T, Lee K, et al. 2015. Role of DNA Methylation in Modulating Transcription Factor Occupancy. Cell Rep 12:1184–1195; doi:10.1016/j.celrep.2015.07.024.

Messerschmidt DM, Knowles BB, Solter D. 2014. DNA methylation dynamics during epigenetic reprogramming in the germline and preimplantation embryos. Genes Dev 28:812–828; doi:10.1101/gad.234294.113.

Nettle D, Frankenhuis WE, Rickard IJ. 2013. The evolution of predictive adaptive responses in human life history. Proceedings Biol Sci 280:20131343; doi:10.1098/rspb.2013.1343.

Pujadas E, Feinberg AP. 2012. Regulated noise in the epigenetic landscape of development and disease. Cell 148:1123–1131; doi:10.1016/j.cell.2012.02.045.

Quilter CR, Cooper WN, Cliffe KM, Skinner BM, Prentice PM, Nelson L, et al. 2014. Impact on offspring methylation patterns of maternal gestational diabetes mellitus and intrauterine growth restraint suggest common genes and pathways linked to subsequent type 2 diabetes risk. FASEB J 28:4868–79; doi:10.1096/fj.14-255240.

Richardson SS, Daniels CR, Gillman MW, Golden J, Kukla R, Kuzawa C, et al. 2014. Society: Don’t blame the mothers. Nature 512:131–132; doi:10.1038/512131a.

Sancho M, Di-Gregorio A, George N, Pozzi S, Sánchez JM, Pernaute B, et al. 2013. Competitive interactions eliminate unfit embryonic stem cells at the onset of differentiation. Dev Cell 26:19–30; doi:10.1016/j.devcel.2013.06.012.

Sharp GC, Salas LA, Monnereau C, Allard C, Yousefi P, Everson TM, et al. 2017. Maternal BMI at the start of pregnancy and offspring epigenome-wide DNA methylation: findings from the pregnancy and childhood epigenetics (PACE) consortium. Hum Mol Genet 26:4067–4085; doi:10.1093/hmg/ddx290.

Slieker RC, Roost MS, van Iperen L, Suchiman HED, Tobi EW, Carlotti FF, et al. 2015. DNA Methylation Landscapes of Human Fetal Development. W. Reik, ed PLoS Genet 11:e1005583; doi:10.1371/journal.pgen.1005583.

Stein Z, Susser M. 1975. Fertility, fecundity, famine: food rations in the dutch famine 1944/5 have a causal relation to fertility, and probably to fecundity. Hum Biol 47: 131–154.

Susser M, Stein Z. 1994. Timing in prenatal nutrition: a reprise of the Dutch Famine Study. Nutr Rev 52: 84–94.

Talens RP, Boomsma DI, Tobi EW, Kremer D, Jukema JW, Willemsen G, et al. 2010. Variation, patterns, and temporal stability of DNA methylation: considerations for epigenetic epidemiology. FASEB J 24:3135–3144; doi:10.1096/fj.09-150490.

Team RDC, R Development Core Team R. 2009. R: A language and environment for statistical computing. R.D.C. Team, ed R Found Stat Comput 1:409; doi:10.1007/978-3-540-74686-7.

Teschendorff AE, Liu X, Caren H, Pollard SM, Beck S, Widschwendter M, et al. 2014. The dynamics of DNA methylation covariation patterns in carcinogenesis. PLoS Comput Biol 10:e1003709; doi:10.1371/journal.pcbi.1003709.

Tobi EW, Goeman JJ, Monajemi R, Gu H, Putter H, Zhang Y, et al. 2014. DNA methylation signatures link prenatal famine exposure to growth and metabolism. NatCommun 5:5592; doi:10.1038/ncomms6592.

Tobi EW, Lumey LH, Talens RP, Kremer D, Putter H, Stein AD, et al. 2009. DNA Methylation differences after exposure to prenatal famine are common and timing-and sex-specific. HumMolGenet 18:4046–4053; doi:10.1093/hmg/ddp353.

Tobi EW, Slagboom PE, van Dongen J, Kremer D, Stein AD, Putter H, et al. 2012. Prenatal famine and genetic variation are independently and additively associated with dna methylation at regulatory loci within IGF2/H19. PLoS One 7:1–11; doi:10.1371/journal.pone.0037933.

Tobi EW, Slieker RC, Stein AD, Suchiman HED, Slagboom PE, van Zwet EW, et al. 2015. Early gestation as the critical time-window for changes in the prenatal environment to affect the adult human blood methylome. IntJEpidemiol 44:1211–1223; doi:10.1093/ije/dyv043.

van den Heuvel J, English S, Uller T. 2016. Disposable Soma Theory and the Evolution of Maternal Effects on Ageing. PLoS One 11:e0145544; doi:10.1371/journal.pone.0145544.

van Iterson M, Tobi EW, Slieker RC, den Hollander W, Luijk R, Slagboom PE, et al. 2014. MethylAid: Visual and interactive quality control of large Illumina 450k data sets. Bioinformatics 30:3435–3437; doi:10.1093/bioinformatics/btu566.

Waggoner MR, Uller T. 2015. Epigenetic determinism in science and society. New Genet Soc 34:177–195; doi:10.1080/14636778.2015.1033052.

Wilcox AJ, Baird DD, Weinberg CR. 1999. Time of implantation of the conceptus and loss of pregnancy. N Engl J Med 340:1796–1799; doi:10.1056/NEJM199906103402304.

Yin Y, Morgunova E, Jolma A, Kaasinen E, Sahu B, Khund-Sayeed S, et al. 2017. Impact of cytosine methylation on DNA binding specificities of human transcription factors. Science 356:eaaj2239; doi:10.1126/science.aaj2239.

